# Inter-subject phase synchronization differentiates neural networks underlying physical pain empathy

**DOI:** 10.1101/841197

**Authors:** Lei Xu, Taylor Bolt, Jason S. Nomi, Jialin Li, Xiaoxiao Zheng, Meina Fu, Keith M. Kendrick, Benjamin Becker, Lucina Q. Uddin

## Abstract

Recent approaches for understanding the neural basis of pain empathy emphasize the dynamic construction of neural networks underlying this multifaceted social cognitive process. Inter-subject phase synchronization (ISPS) is an approach for exploratory analysis of task-based fMRI data that reveals brain networks dynamically synchronized to task-features across participants. We applied ISPS to task-fMRI data assessing vicarious pain empathy in a large sample of healthy participants (n=238). The task employed physical (limb) and affective (faces) painful and corresponding non-painful visual stimuli. ISPS revealed two distinct networks synchronized during physical pain observation, one encompassing anterior insula and midcingulate regions strongly engaged in (vicarious) pain, and another encompassing parietal and inferior frontal regions associated with social cognitive processes which may further modulate and support the physical pain empathic response. No robust network synchronization was observed while processing affective pain, possibly reflecting high inter-individual variation in response to socially transmitted pain experiences. ISPS also revealed networks related to task onset or general processing of physical (limb) or affective (face) stimuli which encompassed networks engaged in object manipulation or face processing, respectively. Together, the ISPS approach permits segregation of networks engaged in different psychological processes, providing additional insight into shared neural mechanisms of empathy for physical pain, but not affective pain, across individuals.

## Introduction

Observing others in pain elicits pain empathetic responses in humans. Empathy, the ability to understand the feelings of others by connecting with those same feelings in one’s self, is a multifaceted social-cognitive process which employs several emotional and cognitive systems, such as affect sharing, simulation, theory of mind and self-other distinction (Shamay-Tsoory, 2011). Empathy for pain refers to vicariously experiencing and – at least to some extent – understanding the feelings of other’s pain. The anterior insula (AI) and the anterior cingulate cortex (ACC) – which represent core nodes of the pain matrix (Price, 2000; Wager et al., 2013) and larger salience network (Uddin, 2015) – respond both during experiencing first-hand pain as well as observing pain in others (Bernhardt & Singer, 2012; Jackson, Rainville, & Decety, 2006; Singer et al., 2004). These brain regions are involved when observing the actual infliction of physical pain in others (e.g. observing that someone cuts his finger with a knife) but also when viewing static or dynamic painful facial expressions of others (Botvinick et al., 2005; Saarela et al., 2006). Meta-analyses of fMRI studies have furthermore confirmed robust engagement of the AI and cingulate regions, specifically dorsal ACC and anterior portions of the midcingulate cortex (MCC) during empathic processes, including pain empathy (Fan, Duncan, de Greck, & Northoff, 2011; Lamm, Decety, & Singer, 2011; Timmers et al., 2018). In addition, the mirror-neuron system which comprises the inferior frontal gyrus (IFG) and inferior parietal lobule (IPL) (Iacoboni & Dapretto, 2006), brain regions related to mentalizing and self-other discrimination such as the temporoparietal junction (TPJ), the medial prefrontal cortex (mPFC), the posterior cingulate cortex (PCC) and the medial temporal lobe (MTL) (Kurczek et al., 2015; Saxe & Kanwisher, 2003; Schurz, Radua, Aichhorn, Richlan, & Perner, 2014; Uddin, Iacoboni, Lange, & Keenan, 2007) are also engaged in pain empathy processing.

Inspired by approaches adapted from network neuroscience, researchers have recently begun to move away from trying to pinpoint specific patterns of regional brain activation associated with pain empathy and to consider pain empathy as a process relying on the dynamic construction of neural networks (Betti & Aglioti, 2016). These network-based approaches have the advantage of moving beyond the cognitive subtraction methodology of identifying functional specialization and can describe the contribution of interactions among brain regions that dynamically associate across time (Kucyi & Davis, 2015). The conventional general linear model (GLM) approach for analysis of task-fMRI data is hypothesis-driven such that a hypothesized reference function that specifies onset and duration of task-specific conditions is convolved with the assumed blood-oxygen-level dependent (BOLD) time courses (Friston et al., 1994). This approach relies on priors with respect to the temporal structure of the task as well as the neural response in terms of the BOLD-signal as described in the hemodynamic response function (HRF). These assumptions may be violated in studies examining complex and dynamic processes like pain empathy, such that behavioral and neural responses to pain stimuli may last longer, and may not immediately stop as soon as the stimuli disappear. Analysis of task-driven brain activity during these paradigms may thus require alternate modeling approaches.

For over a decade, inter-subject synchronization measures of brain activation have been well established in tasks related to auditory stimuli (Hejnar, Kiehl, & Calhoun, 2007), narrated stories (Finn, Corlett, Chen, Bandettini, & Constable, 2018) and movies (Glerean, Salmi, Lahnakoski, Jääskeläinen, & Sams, 2012; Hasson, Nir, Levy, Fuhrmann, & Malach, 2004; Kauppi, Jääskeläinen, Sams, & Tohka, 2010). These studies are based on earlier work demonstrating that brain regions produce similar temporal dynamics across participants experiencing the same task event concurrently (Hanson, Gagliardi, & Hanson, 2009). Recently, an inter-subject phase synchronization (ISPS) analysis which combines the instantaneous phase synchronization measure (Glerean et al., 2012; Kauppi, Pajula, & Tohka, 2014) and independent component analysis (ICA) (Beckmann & Smith, 2004, 2005; Calhoun, Kiehl, & Pearlson, 2008) has been introduced as a means for conducting exploratory analysis of task-based fMRI data (Bolt, Nomi, Vij, Chang, & Uddin, 2018). This approach estimates the task relevant brain networks that dynamically synchronize during the task across participants in a data-driven manner, without dependence on *a priori* reference functions, thus one can potentially gain information about brain responses that is not predicted by the hypothesized temporal structure of the task. There are three methodological and theoretical advantages of the ISPS approach in an exploratory context over the traditional GLM approach: 1) a reference function of task events or HRF is not required to detect task-responsive brain regions, 2) the synchronization approach does not assume that task-driven brain responses follow a single, simple form (e.g. transient, sustained or mixed activation) of voxel-wise activity, and 3) the approach allows for assessing dynamical or time-varying task-related brain networks. To summarize, the ISPS approach is a model-free, data-driven and dynamical measurement of inter-subject synchronization.

The ISPS approach is in practice a form of functional connectivity. However, rather than the more traditional analyses in which functional connectivity is considered a statistical measure quantifying the correlation of time series obtained from *different* brain regions *within the same participant*, ISPS is a measure quantifying the correlation of time series obtained from the *same* brain regions *across multiple participants*. The ISPS approach differs from conventional inter-subject correlation (ISC) analyses in that it is a data-driven, dynamical measure for voxel-wise assessments of synchronization at each time point (Bolt et al., 2018; Glerean et al., 2012; Kauppi et al., 2014). This methodology permits identification of time-varying, task-driven brain network dynamics without dependence on a priori reference functions, and provides a framework for the examination of task-relevant whole-brain temporal dynamics. Compared to another exploratory approach for analysis of task fMRI – the tensor ICA approach (Beckmann & Smith, 2005), the ISPS approach estimates the group-wise synchrony of phase time series rather than the original subject-level BOLD signals, thus revealing unique insights into task-driven brain activity that are not revealed by ICA alone (Bolt et al., 2018). The efficiency and power of the ISPS approach have recently been demonstrated in a simple motor task and a social cognitive task provided by the Human Connectome Project (Barch et al., 2013; Bolt et al., 2018).

In the current study, we applied this data-driven approach to a large sample of fMRI data collected during performance of a pain empathy task employing affective and physical pain empathy stimuli as well as corresponding non-painful control stimuli in order to explore neural network synchronization during vicarious pain empathy.

## Methods

### Participants and task paradigm

252 healthy adults were recruited for the current study. All participants signed written informed consent and received monetary compensation for their participation. The study was approved by the local ethics committee (Institutional Review Board, University of Electronic Science and Technology of China) and was in accordance with the latest revision of the Declaration of Helsinki. Neuroimaging data from six participants were lost due to technical failure. Furthermore, four left-handed participants and four participants with head motion exceeding 3.0 mm translation or 3° rotation were excluded. Consequently, 238 participants (120 males; 17-29 years old, mean age = 21.58 ± 2.32 years) remained in the final analysis. These data have been previously published in a study examining the common and specific associations of autistic traits and alexithymia with neural reactivity (Li et al., 2019). Importantly, the previous study employed a mass-univariate GLM approach, and the focus of the previous study was independent from the aim of the present study.

The pain empathy paradigm (see also Li et al., 2019) employed a blocked design including four experimental conditions (physical pain, affective pain, physical control, affective control). The physical stimuli showed a person’s hand or foot in painful or non-painful everyday situations from a first-person perspective (see Meng et al., 2012) and the affective stimuli consisted of painful and neutral facial expressions from 16 Chinese subjects (8 males) (see Sheng & Han, 2012). A total of 16 picture blocks (4 blocks per condition) were presented in the same pseudorandomized order for all participants and interspersed with a jittered inter-block interval of 8/10/12s showing a red fixation cross, thus permitting the use of the ISPS approach measuring brain synchrony across participants. Each picture block (16s) had four homogeneous stimuli displayed for 3s followed by a 1s white fixation cross on a gray background. The total duration of the task was 436s acquired in a single fMRI run. Participants were required to passively view the stimuli. After scanning, participants were asked to rate the pain intensity and arousal of the stimuli they just viewed in the scanner (see in Table S2 for the rating results).

### Image acquisition and data preprocessing

Neuroimaging data were collected on a 3.0-T GE Discovery MR750 system (General Electric Medical System, Milwaukee, WI, USA). Functional time-series were acquired using a T2*-weighted echo-planar imaging (EPI) sequence (repetition time: 2000 ms; echo time: 30 ms; flip angle: 90°; number of slices: 39 (interleaved ascending order); slice thickness: 3.4 mm; slice gap: 0.6 mm; field of view: 240 × 240 mm^2^; resolution: 64 × 64). To improve normalization of the functional MRI data, high resolution T1-weighted structural images were additionally acquired using a 3D spoiled gradient recalled (SPGR) sequence (repetition time: 6 ms; echo time: minimum; flip angle: 9°; number of slices: 156; slice thickness: 1 mm without gap; field of view: 256 × 256 mm^2^; acquisition matrix: 256 × 256). OptoActive MRI headphones (http://www.optoacoustics.com/) were used to reduce acoustic noise exposure for the participants during MRI data acquisition.

Preprocessing was conducted using standard procedures in SPM12 (Statistical Parametric Mapping, http://www.fil.ion.ucl.ac.uk/spm/), including removal of the first 10 volumes to allow MRI equilibration and active noise cancelling, head motion correction using a six-parameter rigid body algorithm, tissue segmentation and skull-stripped bias-correction for the high-resolution structural images, co-registration of the mean functional image to structural image, normalization (resampling at 3 × 3 × 3 mm) to Montreal Neurological Institute (MNI) space and spatial smoothing with 8mm full-width at half maximum (FWHM) Gaussian kernel. Additionally, we performed denoising using ICA-AROMA (Pruim et al., 2015).

### Inter-subject phase synchronization (ISPS) analysis

Following previous work (Bolt et al., 2018), preprocessing for the synchronization analysis additionally included detrending and filtering (0.01-0.1 Hz) using DPARSF (http://www.restfmri.net/forum/DPARSF, Yan & Zang, 2010). For each subject, time-series of each voxel were extracted and z-transformed. The inter-subject instantaneous phase synchronization analysis (Bolt et al., 2018; Glerean et al., 2012) works by first creating an analytic (i.e. complex-valued) representation of the preprocessed BOLD signal using the Hilbert transformation. We calculated phase synchronization at each time point using a metric known as circular variance. This metric measures the dispersion of phase angles across all subject’s analytic (complex-valued) time series at each time point. This measure provides a single summary statistic across participants at each time point, as opposed to the subject pair-wise average angular distance measure used in previous studies (Bolt et al., 2018; Kauppi et al., 2014). At each time point, we subtracted the circular variance from 1 to obtain the synchronization measure. The synchronization measure varies from 0 to 1, where a value of 1 represents complete similarity of phase signals, and a value of 0 represents the complete absence of similarity of phase signals.

Thus, the result of the instantaneous phase synchronization analysis is a time-series of synchronization values for each voxel in the brain, representing the average synchrony (the average absolute angular distance) across participants for each time point (TR). Rather than a conventional region-of-interest (ROI-based) analysis of synchronization, we chose to use a data-driven independent component analysis (ICA) that incorporates synchronization time signals across the entire brain to estimate possible synchronization networks that appear across the course of the task scan. ICA was implemented through FSL’s MELODIC software (Beckmann, DeLuca, Devlin, & Smith, 2005). This approach is equivalent to a single-subject ICA applied to group-level synchronization time series across all voxels in the brain, rather than the original signal time-courses of all voxels. As in previous work (Bolt et al., 2018), a 10-component ICA solution yielded the highest replicability compared with 15 and 20 component solutions. In the present data set, the 10 component ICA solution was also more replicable compared with the 15 component ICA solution (see Supplementary Materials). Thus, results from the 10-component ICA solution are presented. Components were labeled as unclassified if the spatial weights had characteristic artifact patterns, such as strong weights in white matter, cerebrospinal fluid (CSF) or along the surface of the brain. To further characterized the components, the thresholded (z > 2.3) spatial components were then linked to the well-known Yeo-7 network solution (Yeo et al., 2011) by calculating the percentage of overlapped voxels with each of 7 networks (see Fig. S2). The time course of independent components (ICs) of interest represents the degree of synchronized BOLD activity across all participants at each time point.

### General linear model (GLM) analysis

To compare the brain networks from the phase synchronization analysis to the standard GLM approach, conventional GLM analyses on ICA-AROMA denoised data were conducted using SPM12. On the first level 4 condition-specific regressors (physical pain, affective pain, physical control, affective control) were modelled using a boxcar function and convolved with the canonical HRF. The six head motion parameters were additionally included as nuisance regressors. A 128 s high-pass filter was applied to further control low-frequency noise artifacts. At the second level, one sample *t-*tests were conducted to determine condition-specific activation maps, and physical and affective pain networks respectively, employing subtraction contrasts between the pain and their respective control conditions.

### Association of task block regressors with synchronization time series

To examine the relationship between the synchronization time series resulting from the ICA with each task condition we used the convolved block regressors of the GLM as reference functions. As noted by others (Nummenmaa et al., 2014), conventional double-gamma HRF convolved regressors attempt to model a late undershoot of the HRF, which would not presumably be present in the case of a voxel synchronization time course (which would exhibit no undershoot). Thus, to get the suitable reference functions, we computed another GLM model using the gamma HRF in FSL to account for HRF lag and width, without an undershoot. The association between the synchronization time series and a chosen reference function was computed using the Pearson correlation with Bonferroni correction (8 condition x 9 components, see Table S1). The more positive the correlation, the stronger the association between the reference function and synchronization timeseries from ICs of interest.

## Results

### Conventional GLM results

Brain activation maps produced by the standard GLM approach revealed that all conditions engaged visual cortices and fronto-parietal areas, which may reflect the visual nature of all stimuli and general attention processes (see Fig. 1). Subtraction contrasts between pain and control in both physical and affective conditions revealed typical pain empathy networks. Physical pain compared with the control condition revealed increased activations in bilateral clusters including the IPL, dorsomedial prefrontal cortex (dmPFC), insula and IFG, as well as right lateralized clusters in the MTL, inferior occipital gyrus (IOG), amygdala and thalamus (FWE peak level corrected, *p* < 0.05). Subtraction contrasts between affective pain and control showed increased activation in the bilateral IPL, MTL and TPJ (FWE peak level corrected, *p* < 0.05).

**Figure 1.**
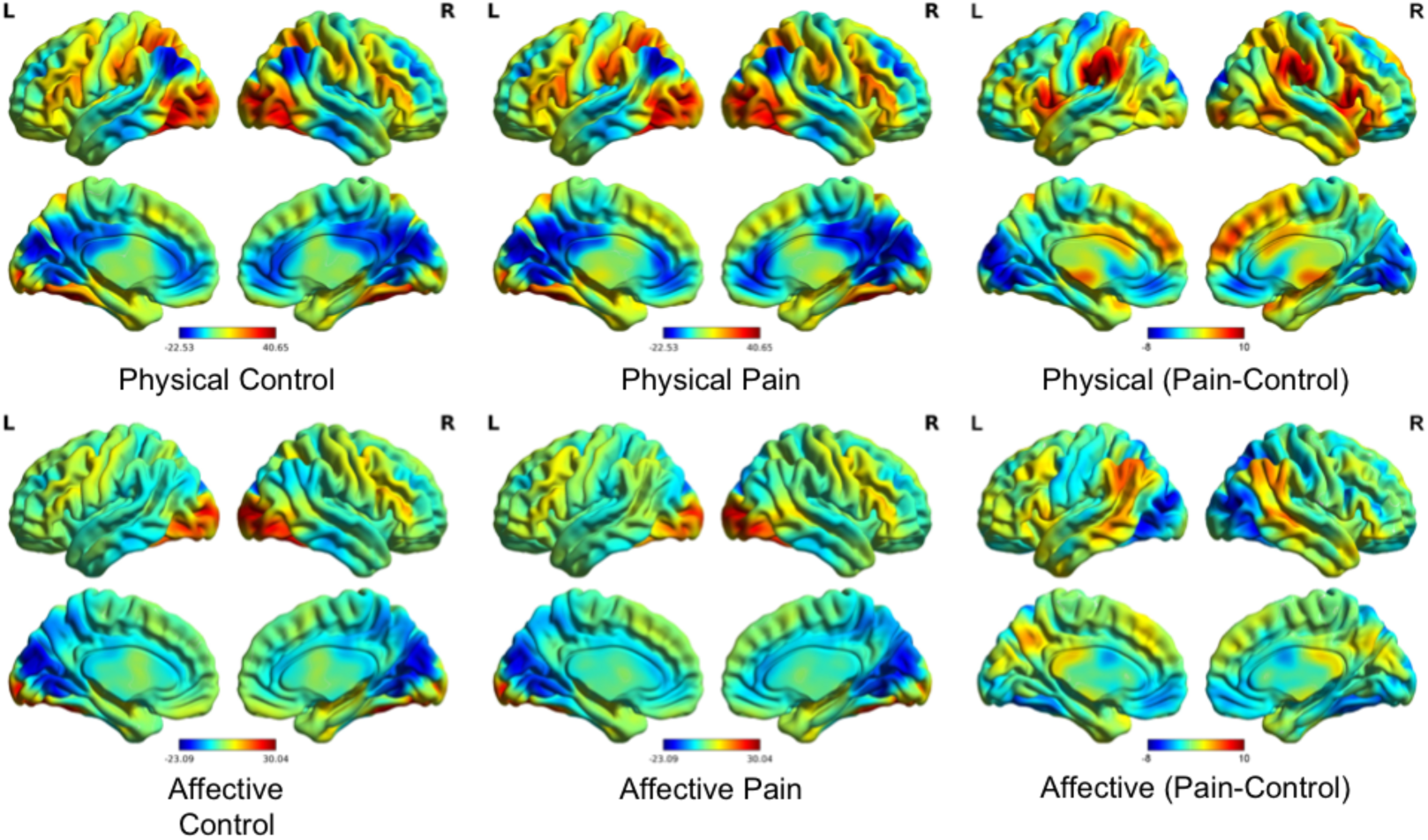
Activation maps (unthresholded) from the conventional GLM approach. The BOLD activation maps for each task condition (middle) as compared to baseline (left), or the relative subtraction contrasts between pain and control in physical and affective conditions, respectively (right).

### Synchronization results

One of the resulting ten ICs was labeled as unclassified because the spatial weights were mostly in white matter and CSF and could not be replicated in the split half sample replication (see Fig. S1). Five of the nine remaining components were related to physical stimuli (domain general networks: C2 and C3; task condition-specific networks: C1, C5, C9) and the other three components corresponded to affective stimuli (domain general networks: C4 and C6; task condition-specific network: C8) and one corresponded to the default mode network (C7) (see Fig. 2).

**Figure 2.**
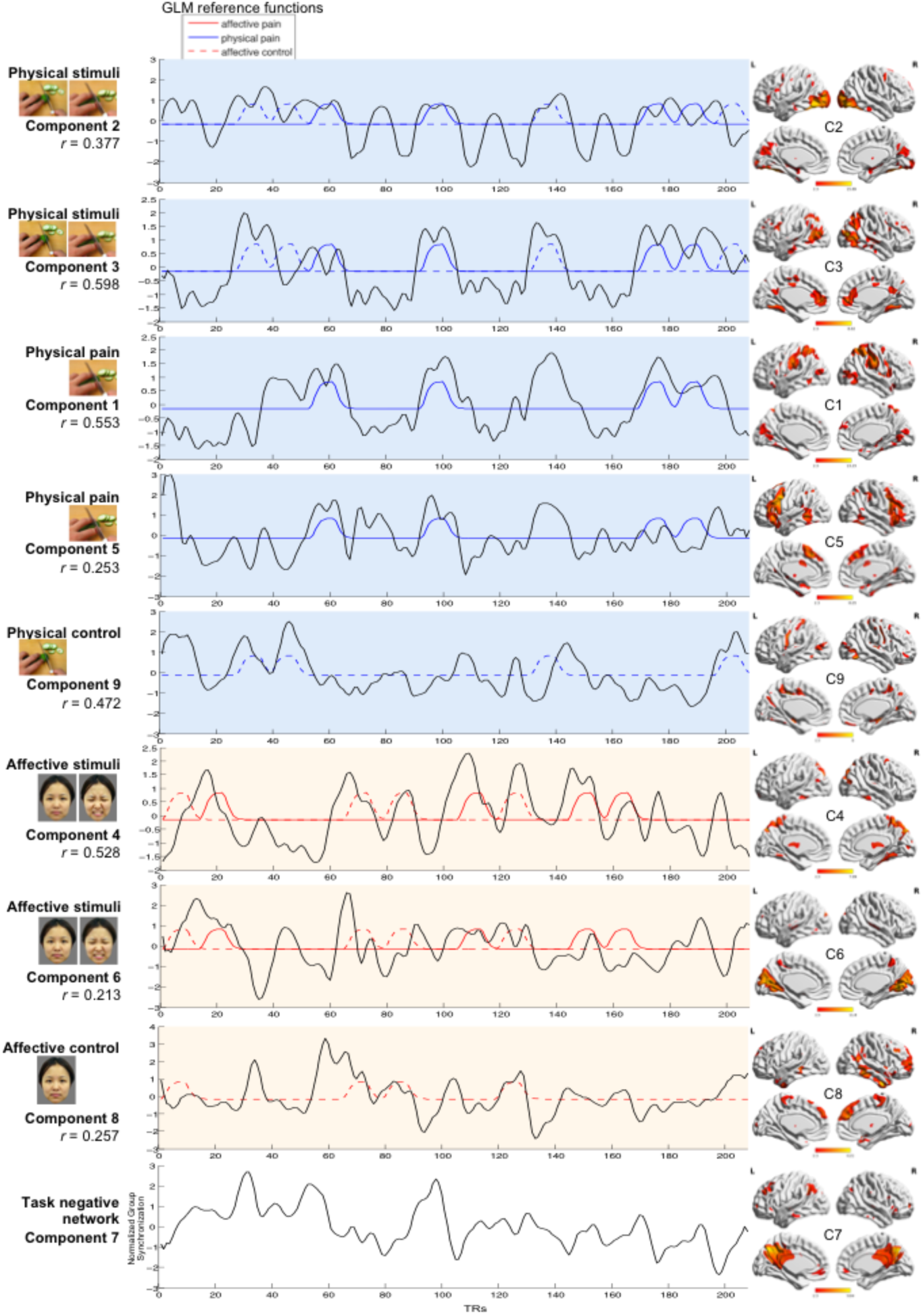
Synchronization of the nine components. (C = Component; R = Right; L = Left). The left panel displays correlation coefficients of each component and the most associated task condition. The middle panal displays the synchronization time courses for each component and the GLM reference functions it is most associated with. The right panel shows the spatial weights for each component (visualized with BrainNet Viewer (Xia, Wang, & He, 2013)).

The synchronization time series of ICA components C2 and C3 were highly correlated with the reference function of physical (limb) stimuli blocks (C2: *r* = 0.377, *p* < 0.001; C3: *r* = 0.598, *p* < 0.001; see Table S1). The spatial map of C2 exhibited the strongest weights in the visual network including the bilateral middle/inferior occipital gyrus, with visual inspection revealing that synchronization peaks of the C2 component were pronounced at the onset of each task block, suggesting that C2 may capture synchronization in a visual network predominantly engaged during the onsets of blocks displaying physical stimuli. In addition, C3 was temporally related to both physical conditions, such that this component significantly correlated with the reference function of physical pain blocks (*r* = 0.423, *p* < 0.001) as well as physical control blocks (*r* = 0.316, *p* < 0.001; Fisher z-test, z = 1.268, *p* = 0.205;; see Table S1), with the spatial pattern of C3 suggesting synchronization in a network incorporating the bilateral middle occipital and middle temporal as well as inferior parietal, and ACC/ventromedial prefrontal cortex (vmPFC) regions. Comparing C3 with the Yeo-7 network solution revealed that C3 included a combination of default mode, visual, dorsal attention and fronto-parietal control networks (see Fig. S2).

The synchronization time series of C1 and C5 were highly correlated with the reference function of physical pain blocks. Although C1 showed a domain general association with all physical stimuli (C1: *r* = 0.568, *p* < 0.001), examination of the condition-specific reference functions revealed that this component demonstrated a stronger association with the temporal reference function of physical pain blocks (*r* = 0.553, *p* < 0.001) rather than physical control blocks (*r* = 0.149, *p* = 0.032; significant difference between the conditions according to Fisher z-test, z = 4.773, *p* < 0.001, Cohen’s *q* = 0.473; medium effect size, details see also Table S1). The spatial pattern of C1 had strongest weights in the inferior/superior parietal regions, including the bilateral postcentral and supramarginal gyrus, as well as additional clusters in the bilateral insula and adjacent IFG. Comparing C1 with the Yeo-7 network solution revealed that C1 had a combination of dorsal attention and visual networks (see Fig. S2). Furthermore, component C5 was also temporally related to the physical pain blocks (*r* = 0.253, *p* < 0.001) rather than physical control blocks (*r* = −0.045, *p* = 0.519; Fisher z-test, z = 3.072, *p* = 0.002, Cohen’s *q* = 0.304; medium effect size). The spatial pattern of C5 had strongest weights in the bilateral AI and the adjacent IFG as well as dmPFC and adjacent MCC regions. Comparing C5 with the Yeo-7 network solution revealed that C5 had a combination of fronto-parietal control and default mode networks (see Fig. S2).

The synchronization time series of C9 were highly correlated with the reference function of physical control blocks (*r* = 0.472, *p* < 0.001) but not physical pain blocks (*r* =−0.344, *p* < 0.001; Fisher z-test, z = 8.821, *p* < 0.001, Cohen’s *q* = 0.871; large effect size; see Table S1) with the spatial pattern indicating strongest weights in the bilateral precentral and postcentral gyrus, which involve the somatomotor network.

The synchronization time series of C4 and C6 were highly correlated with the reference function of affective (face) stimuli blocks (C4: *r* = 0.528, *p* < 0.001; C6: *r* = 0.213, *p* = 0.002; see Table S1). The spatial pattern of C6 had strong weights predominantly located in the medial and superior occipital visual regions including the cuneus, calcarine and lingual gyrus, and regions engaged in social and face processing including the fusiform and superior temporal gyrus. And precuneus. Thus, C6 may capture visual network related to face processing. C4 was temporally related to both affective pain blocks (*r* = 0.396, *p* < 0.001) and affective control blocks (*r* = 0.255, *p* < 0.001; Fisher z-test, z = 1.600, *p* = 0.110; see Table S1) and had strong spatial weights in the bilateral posterior cerebellum, occipital and temporal regions including cuneus, fusiform gyrus and precuneus, as well as the thalamus and superior frontal regions. Comparing C4 with the Yeo-7 network solution revealed that C4 had a combination of dorsal attention and visual networks (see Fig. S2).

The synchronization time series of C8 was correlated with the reference function of affective control blocks (neutral facial expression, *r* = 0.257, *p* < 0.001) but not affective pain blocks (*r* =−0.132, *p* < 0.001; Fisher z-test, z = 3.996, *p* < 0.001, Cohen’s *q* = 0.395; large effect size; see Table S1) and the spatial pattern indicated strongest weights in the right inferior and middle temporal gyrus including the fusiform gyrus, and superior mPFC. Comparing C8 with the Yeo-7 network solution revealed that C8 had a combination of default mode, fronto-parietal control and limbic networks (see Fig. S2). None of the components specifically related to the temporal reference function of the affective pain condition.

Finally, the synchronization time series of C7 was not specifically associated with a specific task condition (all *r* < 0.133, *p* > 0.055) and was observed to have strong spatial weights predominantly located in the default mode network (precuneus, posterior cingulate cortex and vmPFC).

Quantitatively comparing each component to the Yeo-7 network solution (Yeo et al., 2011) by calculating the percentage of overlapped voxels confirmed the results from the visual inspection of the overlap (see Fig. S2 for details).

## Discussion

The present study employed a data-driven ISPS approach to a large task-fMRI data set of healthy participants that used visual stimuli to engage pain empathic brain networks by presenting affective and physical pain stimuli as well as corresponding non-painful control stimuli. The synchronization approach determined networks that were engaged across processing of physical or affective stimuli, respectively. Moreover, task condition-specific networks were observed for physical pain, physical control and affective control stimuli, while no robust networks were determined for the affective pain stimuli.

With respect to the processing of physical stimuli, the ISPS approach reliably identified networks engaged in domain-general processing of physical stimuli as well as pain empathy specific networks. Components C2 and C3 were associated with the reference functions modelling both physical pain and physical control conditions, suggesting general inter-subject synchronization of these components for physical stimuli irrespective of pain empathic processing. Component C2 predominately captured a network encompassing primary visual processing areas in the medial and inferior occipital lobe, likely reflecting processing of low-level visual features and object categorization (DiCarlo, Zoccolan, & Rust, 2012). Visual inspection revealed pronounced synchronization during the beginning of physical stimuli blocks, which may reflect stronger engagement of object categorization or novelty detection at the onset of the condition-specific block (Ranganath & Rainer, 2003) or unspecific mechanisms related to repeated presentation of similar visual stimuli in visual processing areas such as habituation or repetition suppression processes (Vidyasagar, Stancak, & Parkes, 2010). Component C3 primarily encompassed middle temporal and inferior parietal, as well as mPFC regions. This synchronized network overlaps with parietal and temporal regions engaged during action observation (Caspers, Zilles, Laird, & Eickhoff, 2010; Molenberghs, Cunnington, & Mattingley, 2012), including observation of complex hand-object manipulations (Errante & Fogassi, 2019) that have been determined employing traditional BOLD level subtraction methods. The mPFC is a functionally highly heterogenous region involved in a broad range of emotional and cognitive functions, which in concert with parietal and temporal regions, supports social cognitive functions including mentalizing during action observation and decoding of goals based on observed body-part motions (Spunt, Satpute, & Lieberman, 2010; Van Overwalle & Baetens, 2009).

Consistent with previous studies employing hypothesis driven GLM approaches and subtraction contrasts comparing physical pain with matched non-painful stimuli, the synchronization time series of two components (C1, C5) specifically correlated with the temporal reference function of physical pain blocks. The components encompassed a network primarily including the inferior and lateral parietal regions, the postcentral and supramarginal gyrus, AI and adjacent IFG as well as the dmPFC and adjacent MCC regions. Previous meta-analytic results from studies employing subtraction contrasts revealed a highly overlapping network engaged during empathic responses, including pain empathy (Fan et al., 2011; Lamm et al., 2011; Timmers et al., 2018). More specifically, component C5 exhibited predominately associations with core nodes of the network engaged in experiencing first-hand as well as vicarious pain, such as the AI and MCC (Bernhardt & Singer, 2012; Jackson et al., 2006; Singer et al., 2004), whereas component C1 encompassed regions engaged in social cognitive processes that coactivate with empathy responses (Bernhardt & Singer, 2012; Shamay-Tsoory, 2011) and may support or modulate the experience of empathy, including the inferior parietal, inferior frontal and dmPFC regions engaged in mirror neuron(Molenberghs et al., 2012) and mentalizing processes (Schurz et al., 2014). Together this suggests that the ISPS approach can identify the core networks engaged in pain empathic processes and, in addition, compared with the standard subtraction method, it can differentiate networks primary engaged in the pain-associated response from social cognitive networks considered to modulate and support the pain empathic response. The physical control condition was additionally associated with a separate component (C9) primarily encompassing bilateral precentral and postcentral gyrus.

Two components (C4, C6) were correlated with the reference function for both affective pain and affective control (facial) stimuli, suggesting that these components may capture aspects underlying general face processing independent of pain. Component C6 encompassed primary visual processing regions in the occipital cortex, including medial occipital and calcarine regions, possibly reflecting primary visual processing of facial stimuli. C4 additionally encompassed temporal and superior frontal regions strongly involved in face processing and face recognition, such as the fusiform gyrus (Fusar-Poli et al., 2009; Sabatinelli et al., 2011), suggesting that the synchronization approach was able to differentiate the networks engaged in these sub-processes. The affective control condition was additionally associated with a separate component (C8) primarily encompassing inferior temporal and superior mPFC regions.

Surprisingly, the reference function for affective pain stimuli was not significantly associated with any of the identified components, suggesting that the concomitant pain information may have interfered with synchronicity across participants. Although the present standard GLM subtraction analysis and previous meta-analysis encompassing studies using facial pain stimuli revealed activation in pain empathy networks (Timmers et al., 2018) a corresponding network that specifically associated with the reference function of the affective pain blocks was not found using the synchronization approach. This might be explained in the context of the group level strategy the ISPS is based upon, such that larger individual differences during affective pain observation may have contributed to the lack of synchronized networks in this condition. If the affective pain stimuli induced highly variable responses across participants, this inter-subject variability would reduce inter-subject synchronization during this condition. Previous studies suggest that dimensional variables such as trait alexithymia and autism (Li et al., 2019) as well as categorical variables including sex and genotype (Warrier et al., 2018) modulate processing of painful faces. Furthermore, as a highly intense negative emotion, the painful facial expression is very similar to the expression of highly intense positive emotions, such as orgasm and victory (Aviezer, Trope, & Todorov, 2012; Hughes & Nicholson, 2008). Thus, the affective pain pictures can be considered as the most ambiguous in the current paradigm – the other stimuli are pretty easily identifiable to participants, however the painful faces without any context (eg. displaying a noxious agent in difference to the physical pain pictures) may induce widely varying interpretations as a function of the previous experience of the participants. In support of this interpretation, we found that post-scanning subjective ratings by the participants revealed that the standard deviation of both pain intensity ratings (SD = 21.43) and arousal ratings (SD = 20.13) for affective pain stimuli were higher than these for physical pain stimuli (pain intensity: SD = 17.18; arousal: SD = 17.53; see Table S2). This further confirms that affective pain perception has higher variability across participants in terms of subjective experience. Finally, a component encompassing core regions of the default mode network, specifically posterior parietal and medial frontal regions (Andrews-Hanna, Reidler, Sepulcre, Poulin, & Buckner, 2010) did not synchronize with any of the task-dependent reference functions.

The findings of the present study need to be considered in the context of some limitations. First, despite the split half replication approach employed here, replications in independent samples are required to fully elucidate the robustness of the findings. Second, the ISPS approach provides a flexible exploratory data-driven approach for task-fMRI data to identify the common and homogenous task/stimuli related networks across participants, however it may be not suitable for tasks which have randomized/counterbalanced presentation order or stimuli which induce variable or non-homogeneous responses across participants (eg. affective pain stimuli here), or when aiming to differentiate individual differences such as sex differences.

Together, the present results further demonstrate that the ISPS approach may represent a valuable exploratory analysis method that can reveal network synchronization in the context of task-based fMRI analyses and can separate networks that support complex social emotional processes such as empathy, especially empathy for physical pain but not affective pain. In the context of growing evidence for dysregulations in pain empathic processes in participants with high levels of pathology-relevant traits such as alexithymia or autism (Bird & Viding, 2014; Li et al., 2019) as well as in patient populations with depression (Xu et al., 2019) or schizophrenia (Vistoli, Lavoie, Sutliff, Jackson, & Achim, 2017) the ISPS method may furthermore permit dissection of impaired network-level integration underlying social cognitive deficits.

## Acknowledgements

This work was supported by the National Key Research and Development Program of China (Grant No. 2018YFA0701400), the National Natural Science Foundation of China (NSFC, 91632117 to BB; 31530032 to KK), the Sichuan Science and Technology Department (2018JY0001 to BB), National Institute of Mental Health award (R01MH107549 to LQU) and a grant from the Canadian Institute for Advanced Research (to LQU). The authors report no biomedical financial interests or potential conflicts of interest.

## Supplementary Material

### Assessment of number of independent components and replication

We compared 10 and 15 component ICA solutions in the current data set. All participants were randomly split into two subgroups (original sample and confirmatory sample) three times for replication and assessment of number of components. ‘Replicability’ was measured in three ways: 1) the number of components that had ‘matching’ component pairs in the confirmatory sample in terms of visual examination between the original and confirmatory ICA solutions, 2) the spatial correlation between the ‘matching’ component pairs from the original and confirmatory ICA solutions, and 3) the temporal correlation between the ‘matched’ components from the original and confirmatory ICA solutions. This replicability analysis also allowed an assessment of the degree to which the synchronization approach replicates across samples. Results suggested that the 10-component ICA solution (1st/2nd/3rd time of grouping: matching components: 9/9/9 out of 10; rSpatial: 0.69/0.66/0.64, rTemporal: 0.73/0.70/0.69) was more replicable compared with the 15 component ICA solution (matching components: 13/13/13 out of 15; rSpatial: 0.54/0.61/0.52, rTemporal: 0.65/0.67/0.63). Thus, results from the 10-component ICA solution are presented.

**Fig. S1.**
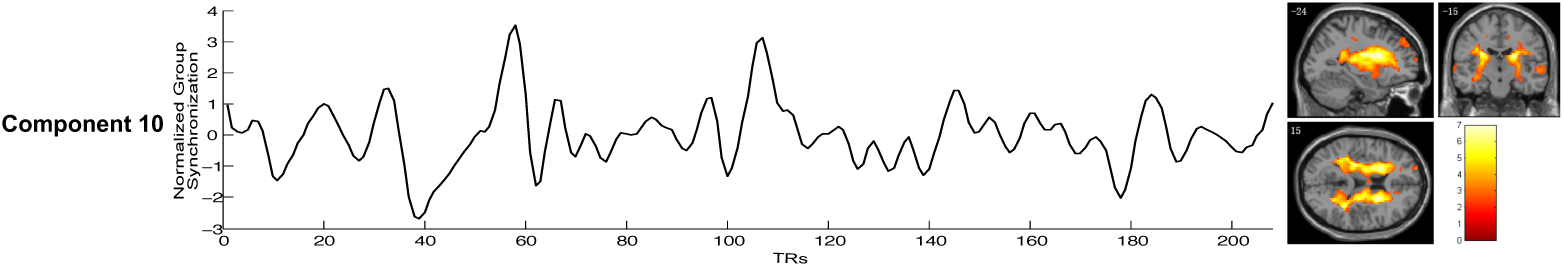
Unclassified component in pain empathy task. C10 was labeled as unclassified based on voxel weight spatial patterns in white matter and cerebrospinal fluid regions. This component also could not be replicated in the split half sample replication.

**Fig. S2.**
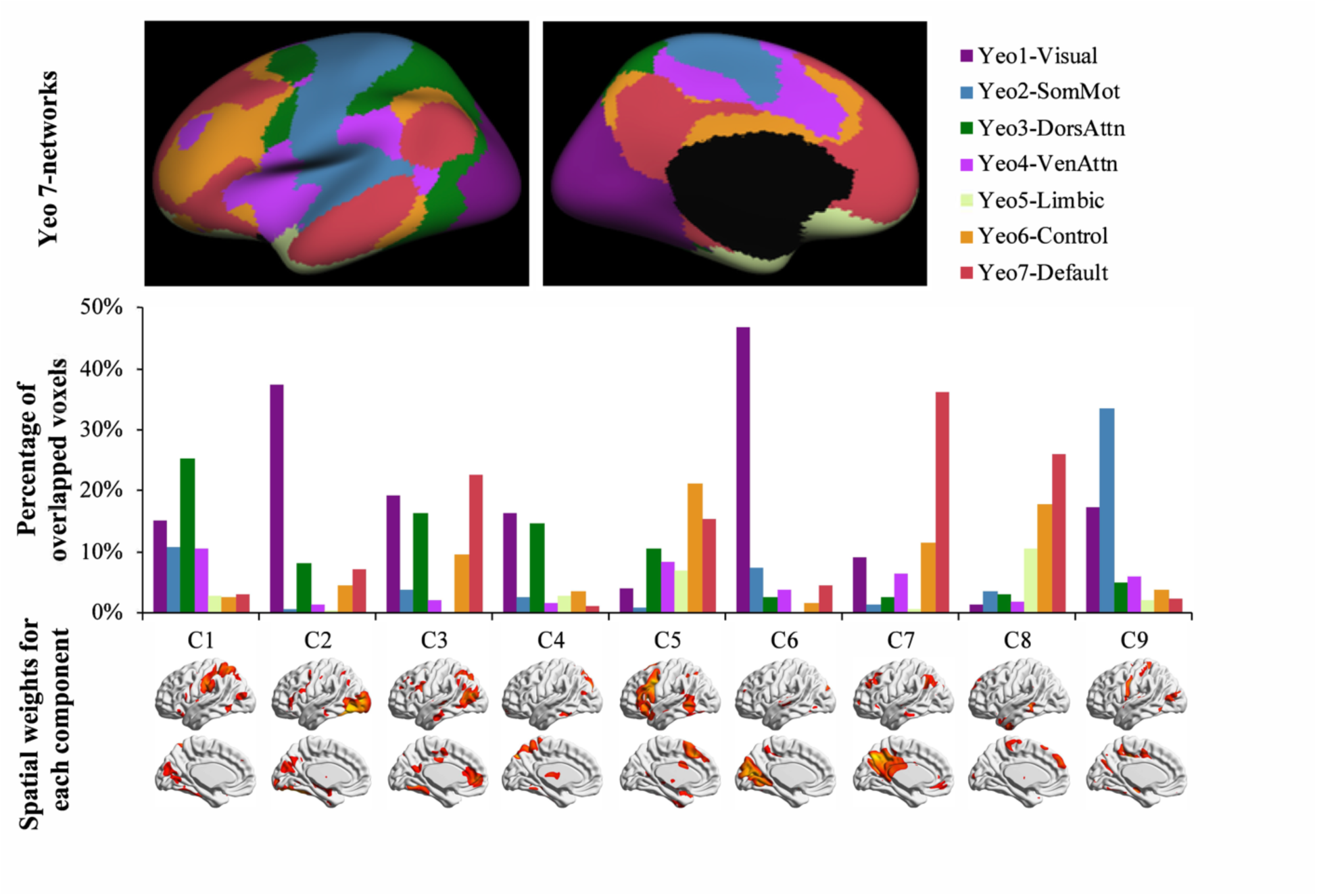
Overlap with 7 Yeo Networks for each component.

**Table S1.**
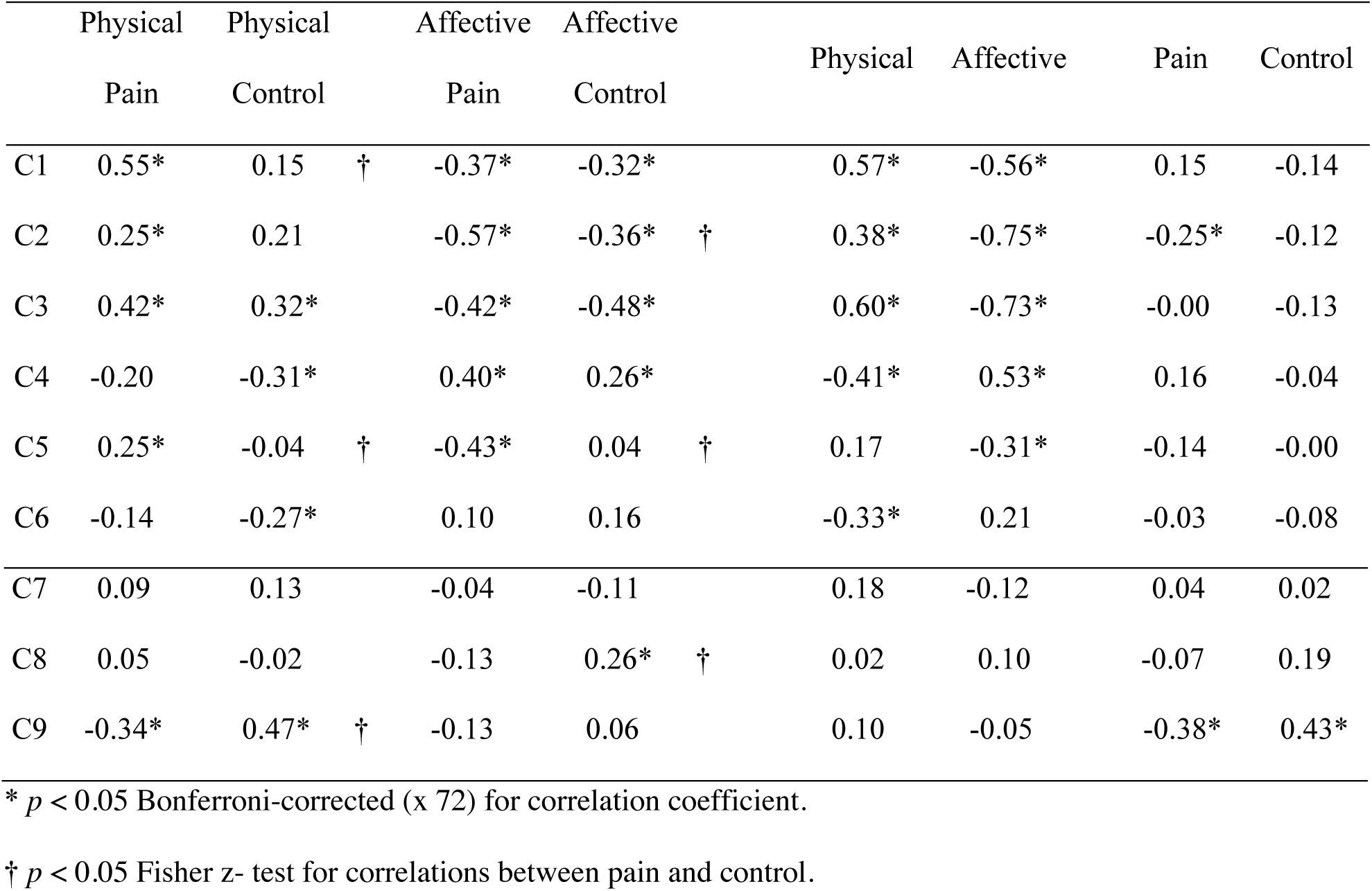
Temporal correlation between ICA components and GLM hypothesized reference functions.

**Table S2.** Subjective ratings for stimuli (*M* ± *SD*)

| Ratings | Categories of stimuli |  |  |  |
| --- | --- | --- | --- | --- |
|  | affective control | physical control | affective pain | physical pain |
| Pain intensity | 5.09 ± 7.34 | 8.49 ± 9.68 | 53.64 ± 21.43 | 71.43 ± 17.18 |
| Arousal | 16.79 ± 18.72 | 22.23 ± 18.74 | 54.48 ± 20.13 | 72.57 ± 17.53 |

